# Epigenetic gene-expression links heart failure to memory impairment

**DOI:** 10.1101/2020.01.22.915637

**Authors:** Rezaul Islam, Dawid Lbik, Sadman Sakib, Raoul Maximilian Hofmann, Tea Berulava, Martí Jiménez Mausbach, Julia Cha, Elerdashvili Vakhtang, Christian Schiffmann, Anke Zieseniss, Dörthe Magdalena Katschinski, Farahnaz Sananbenesi, Karl Toischer, Andre Fischer

**Author notes:** equal contribution (Dawid Lbik, Sadman Sakib). equal contribution (Karl Toischer, Andre Fischer).

## Abstract

In current clinical practice care of diseased patients is often restricted to separated disciplines. However, such an organ-centered approach is not always suitable. For example, cognitive dysfunction is a severe burden in heart failure patients. Moreover, these patients have an increased risk for age-associated dementias. The underlying molecular mechanisms are presently unknown and thus corresponding therapeutic strategies to improve cognition in heart failure patients are missing. Using mice as model organisms we show that heart failure leads to specific changes in hippocampal gene-expression, a brain region intimately linked to cognition. These changes reflect increased cellular stress pathways which eventually lead to loss of neuronal euchromatin and reduced expression of a hippocampal gene cluster essential for cognition. Consequently, mice suffering from heart failure exhibit impaired memory function. These pathological changes are ameliorated via the administration of a drug that promotes neuronal euchromatin formation. Our study provides first insight to the molecular processes by which heart failure contributes to neuronal dysfunction and point to novel therapeutic avenues to treat cognitive defects in heart failure patients.

## Introduction

Traditionally, clinical medicine is organized by organ-centered disciplines which is reflected in the currently applied diagnostics and treatments of patients. This approach has been also commonly adopted in research strategies but it is becoming evident that novel interdisciplinary efforts are needed to improve therapies of complex diseases. For example Heart failure (HF) is a complex, debilitating condition afflicting millions of people worldwide (Savarese & Lund, 2017). However, in addition to the detrimental phenotypes linked directly to cardiac dysfunction, cognitive deficits present a major burden to patients with HF (Ampadu & Morley, 2015, Hajduk, Lemon et al., 2013, Pressler, Subramanian et al., 2010) (Doehner, Ural et al., 2017). Moreover, epidemiological studies have clearly demonstrated that HF significantly increases the risk for dementia and age-associated neurodegenerative diseases such as Alzheimer’s disease (AD) (Angermann, Frey et al., 2012, Cermakova, Lund et al., 2015, Satizabal, Beiser et al., 2016). In line with these observations, a consistent finding in HF patients is a substantially reduced cerebral blood flow (Roy, Woo et al., 2017) and imaging studies reveal subsequent structural and functional cerebral alterations including changes in key regions linked to memory formation, such as the hippocampus (Kumar, Woo et al., 2011) (Pan, Kumar et al., 2013) (Kumar, Yadav et al., 2015) (Woo, Ogren et al., 2015). However, how HF affects hippocampal function at the molecular level remains to be explored and thus effective therapies to manage cognitive impairment if HF patients do not exist yet. On the contrary, the therapeutic approaches currently used to treat cardiac phenotypes in HF patients lack evidence for improving cognition (Cleland, Daubert et al., 2005) (Arnold, Liu et al., 2006, Frigerio & Roubina, 2005) or have even been linked to an increased incidence of AD (Pressler et al., 2010) (Khachaturian, Zandi et al., 2006) (Galli & Lombardi, 2014) (Solomon, Rizkala et al., 2017), suggesting that HF may lead to long-lasting adaptive changes in neurons that can persist despite improvement of cardiac function. Thus, a better understanding of HF-mediated molecular alterations in neurons is of utmost importance but corresponding data is lacking. Consequently, international organizations such as the European Society of Cardiology (ESC) have recommended that cardiology and dementia research experts should team-up to identify therapeutic interventional options for managing cognitive impairment in subjects with HF (Ponikowski, Voors et al., 2018). In this study, we took on this challenge and show that heart failure leads to specific changes in hippocampal gene expression that are linked to memory impairment. Targeting aberrant gene expression via epigenetic drugs ameliorates these phenotypes suggesting a key role of this process in HF-mediated cognitive dysfunction. Moreover, our data suggest that therapeutic strategies directed towards epigenetic gene-expression provide a therapeutic avenue to improve cognition in HF patients and ameliorate their risk to develop AD.

## Results

### Heart failure in CamkIIδc TG mice leads to hippocampal gene expression changes indicative of dementia

With the aim to elucidate the molecular processes by which cardiovascular dysfunction leads to memory impairment and an increases the risk for dementia, we decided to employ a well-established mouse model for heart failure in which cardiomyocyte-specific kinase CamkIIδc is overexpressed under the control of the alpha-MHC promoter (CamkIIδc TG mice) (Maier, Zhang et al., 2003). Thus, overexpression of CamkIIδc is specific to cardiomyocytes and is not detected in other organs, including the brain (Maier et al., 2003), making it a *bona fide* model to study the impact of heart failure on brain function (Fig 1A). We reasoned that this well-defined genetic heart failure model would be superior to other experimental approaches linked for example to cerebral hypoperfusion such as carotid artery occlusion, since it allowed us to study brain function in response to the very precise and exclusive manipulation of cardiac tissue. In line with previous findings, 3-month-old CamkIIδc TG mice displayed heart failure with left ventricular dilatation, impaired ejection fraction, and increased heart mass (Fig 1B, C), whereas the overall body weight was not affected (*P* = 0.863 for CamkIIδc TG vs control mice, n =8, unpaired *t*-test). As a first approach to study the impact of cardiac dysfunction on brain plasticity we decided to analyze the transcriptome of the hippocampal CA1 region in 3-month-old CamkIIδc TG mice (Fig 1D). This was based on data showing that (1) gene expression is a sensitive molecular correlate of memory function and is de-regulated in dementia patients and corresponding mouse models (Fischer, 2014a); (2) the hippocampal CA1 regions is essential for spatial reference memory in rodents and humans and is affected early in AD (Fischer, 2014a) and (3) imaging data show functional changes of the hippocampal CA1 region in patients with HF (Woo et al., 2015). RNA-seq analysis revealed substantial changes in the CA1 transcriptome of 3-month old CamkIIδc TG and control mice that were obvious in a principle component analysis (PCA; Fig 1D). Namely, 1780 genes were up-regulated and 2014 genes were down-regulated in CamkIIδc TG when compared to the control group (Fig. 1E; supplemental table 1). Comparison of the differentially expressed genes to previously reported cell-type specific gene expression datasets (Merienne, Meunier et al., 2019) revealed that up-regulated genes were linked to neurons, microglia and astrocytes, while down-regulated genes were mainly associated with neurons (Fig 1F). Further pathway analysis showed that up-regulated genes are related to cellular stress response pathways such as oxidative and endoplasmic reticulum (ER) stress (Fig. 1G, supplemental table 1), while down-regulated genes are linked to cognition, protein folding and processes related to protein methylation (Fig 1G, supplemental table 1). We decided to confirm the RNA-sequencing data by testing differential expression for selected genes representing changes related to increased cellular stress processes, in this case “ER stress” and down-regulated processes such as “protein methylation”. qPCR analysis confirmed increased expression of the ER stress-related genes Fez1, Fez2 and Bcap31 (Fig 1H). We also tested the expression of several histone 3 lysine 4 (H3K4) specific lysine methyltransferases (Kmts), since these pathways were detected in the RNA-seq data and several of the Kmt’s, such as Kmt2a, were found to be essential for memory formation (Gupta, Kim et al., 2010, Kerimoglu, Agis-Balboa et al., 2013) (Jakovcevski, Ruan et al., 2015) (Kerimoglu, Sakib et al., 2017). Indeed, we observed that Kmt2a and Kmt2d were significantly down-regulated in CamkIIδc TG mice (Fig 1H). Specificity of this observation was demonstrated by the fact that other H3K4 methyltransferases such as Kmt2b and Kmt2c were not differentially expressed.

**Fig. 1:**
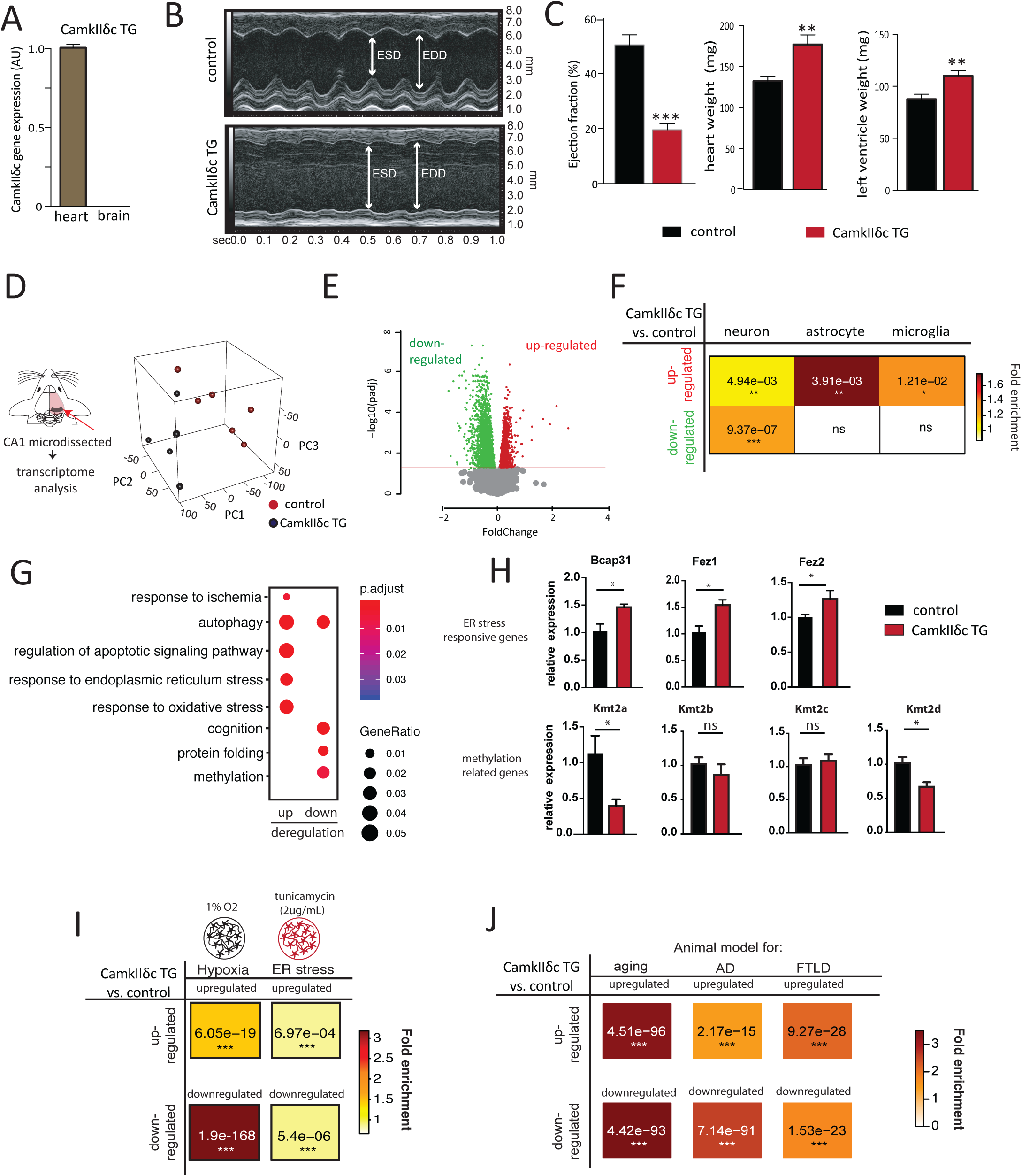
Heart failure in CamkIIδc TG mice is linked to aberrant hippocampal gene-expression. **A.** qPCR data showing expression of the CamkIIδc in the brain and heart of 3 month old CamkIIδc TG mice; n=4/group; \**P* <0.05). **B.** Representative M-mode images from left ventricle from CamkIIδc TG and control mice. ESD: Left ventricle end systolic diameter, EDD: Left ventricle diastolic diameter. **C.** Left panel: Ejection fraction is significantly decreased in CamkIIδc TG mice (n =8) when compared to control mice (n=5; \**P* <0.05). Heart weight (middle panel) and left ventricle weight (right panel) are increased in CamkIIδc TG (n=8) compared to control (n=5; \**P* <0.05). **D.** Experimental scheme for RNA-seq analysis that was performed from hippocampal CA1 region of CamkIIδc TG mice (n=6) and control mice (n=5) at 3 month of age. Right panel shows principle component analysis (PCA) of the gene-expression data. The first principle component (PC1) can explain 42 % of the variation between two groups. **E.** Volcano plot showing differentially expressed genes (FDR<0.05). Red color indicates up-regulation while blue represents down-regulation of transcripts. **F.** Hypergeometric overlap analysis comparing genes deregulated in CamkIIδc TG mice to genes uniquely expressed in neurons, astrocytes or microglia. **G.** Dot plot showing Top GO biological processes after removing redundant GO terms using Rivago. **H.** qPCR quantification of selected genes reflecting ER-stress or protein methylation-related processes (n = 5/group) *p<0.05, Unpaired t-test; two-tailed Data is normalized to Hprt1 expression. **I.** Hypergeometric overlap analysis comparing genes deregulated in CamkIIδc TG mice to genes deregulated under hypoxia conditions and in response to tunicamycin-induced ER-stress **J.** Hypergeometric overlap analysis comparing genes deregulated in CamkIIδc TG mice to genes deregulated in hippocampal tissue from animal models of memory impairment and neurodegeneration. *p<0.05, **p<0.01, ***p<0.001Unpaired t-test; two-tailed. Error bars indicate SEM.

The observation that genes implicated with oxidative and ER stress are increased in the hippocampus of CamkIIδc TG mice is in line with previous findings linking heart failure to hypoxia as a consequence of cerebral hypoperfusion (Bikkina, Levy et al., 1994) (Verdecchia, Porcellati et al., 2001) (Perlman, 2007). The concomitant down-regulation of genes linked to cognition let us to hypothesize about a potential link between the observed cellular stress-related gene-expression changes and the decreased expression of genes associated with cognition. Namely, we wondered if the decreased expression of genes linked to cognition could be a consequence of the activation of cellular stress pathways. We decided to test this hypothesis further with a focus on hypoxia and ER-stress as key cellular stress pathways. Since data on the effects of hypoxia and ER-stress on hippocampal gene-expression at the genome-wide level is still rare, we decided to performed RNA-sequencing from mixed hippocampal neuronal cultures that were subjected to either hypoxia or ER-stress. First, we analyzed hypoxia. Differential expression analysis revealed a substantial amount of genes that were differentially expressed in response to hypoxic conditions (supplemental table 2). We then compared the genes up-and down-regulated in hippocampal cultures in response to hypoxia to the genes up- and down-regulated in the hippocampus of CamkIIδc TG mice. This analysis revealed a significant overlap of not only up - but also down-regulated genes suggesting that hypoxic conditions are sufficient to induce gene-expression changes similar to that detected in the hippocampus of mice suffering from HF (Fig 1I). We employed the same experimental settings to test the impact of ER-stress that can be modeled via the administration of tunicamycin. Thus, RNA-sequencing was performed from mixed hippocampal neuronal cultures upon treatment with tunicamycin (Supplemental table 3). Our data show that genes de-regulated in response to tunicamycin also significantly overlap with genes affected in CamkIIδc TG mice, although to a lesser extend when compared to hypoxia (Fig 1I). In sum, these data suggest a scenario in which heart failure that is linked to cerebral hypoperfusion leads to hypoxia, oxidative and ER-stress related hippocampal gene-expression changes which are upstream of the reduced expression of neuronal genes important for cognition. Taken into account that impaired expression of genes essential for cognitive function is also a key hallmark of dementia, these data provide a plausible hypothesis to explain - at least in part – cognitive dysfunction in response to HF. To provide further evidence for this hypothesis, we first retrieved published datasets in which brain-specific gene-expression changes were reported in mouse models with impaired memory function, namely models for aging-associated memory decline (Benito, Urbanke et al., 2015), models for AD (Gjoneska, Pfenning et al., 2015) and Fronto-temporal dementia (FTLD) (Swarup, Hinz et al., 2018). We compared these datasets to the transcriptional alterations observed in the hippocampus of CamkIIδc TG mice (Fig 1J). Interestingly, there was a significant overlap of genes up-regulated in the hippocampus of CamkIIδc TG mice and genes up-regulated in the hippocampus of cognitively impaired old mice, in CK-p25 mice representing a model for AD-like neurodegeneration and in the cortex of FVB mice, representing a mouse model for fronto-temporal dementia FTLD (Fig 1J). Similarly, genes down-regulated in the hippocampus of CamkIIδc TG mice significantly overlapped with the genes down-regulated in models for aging, AD-like neurodegeneration and FTLD (Fig 1J).

Thus, the hippocampal gene-expression signature observed in response to heart failure partly overlaps to the gene-expression changes detected in cognitive diseases. On this basis we hypothesized that aberrant hippocampal gene-expression and especially the decreased expression of learning and memory genes could be a central process in heart failure mediated cognitive impairment and might therefore represent a suitable target for therapeutic intervention. To further substantialize and test this hypothesis, we first decided to first analyze memory function in CamkIIδc TG mice directly.

### Heart failure in CamkIIδc TG is associated with impaired hippocampus-dependent memory consolidation

Three-month-old CamkIIδc TG (n=16) and control mice (n=13) were subjected to behavioral testing. Importantly, when subjected to the open field test, CamkIIδc TG and control mice traveled similar distances with the same speed, indicating that explorative behavior and basal motor-function is normal in CamkIIδc TG mice (Fig 2A). Both groups also spent similar time in the center of the open field arena, suggesting that anxiety behavior is not affected in CamkIIδc TG mice (Fig 2A). Subsequently, mice were subjected to the Barnes Maze, a hippocampus-dependent spatial navigation-learning test (see methods for details). Two-way ANOVA analysis revealed that CamkIIδc TG mice spent significantly more time to find the escape hole when compared to littermate controls (Fig 2B). These data suggest that hippocampus-dependent memory function is impaired in CamkIIδc TG mice. A detailed analysis of the different strategies to find the escape hole confirmed this observation and revealed that in comparison to control mice, CamkIIδc TG mice failed to adapt hippocampus-dependent strategies (direct, short and long chaining approaches), which are generally considered to depend on higher cognitive abilities than the other strategies (Fig. 2C). To quantify this observation, we calculated the cumulative strategy score (see methods for details) that was significantly reduced in CamkIIδc TG mice when compared to the control group (Fig 2D), further confirming that CamkIIδc TG mice exhibit impaired hippocampus-dependent learning abilities. We also assayed memory retrieval 24h after the final day of training by placing the mice into the Barnes Maze arena with the escape hole being closed and measured the visits to the escape hole. The number of visits to the escape hole during the 120 sec test period was significantly lower in CamkIIδc TG mice when compared to the control group, indicating impaired retrieval of spatial memories (Fig 2E). In summary, these findings are in line with our gene-expression data (See Fig. 1) and show that CamkIIδc overexpression-induced heart failure leads to cognitive deficits.

**Fig. 2:**
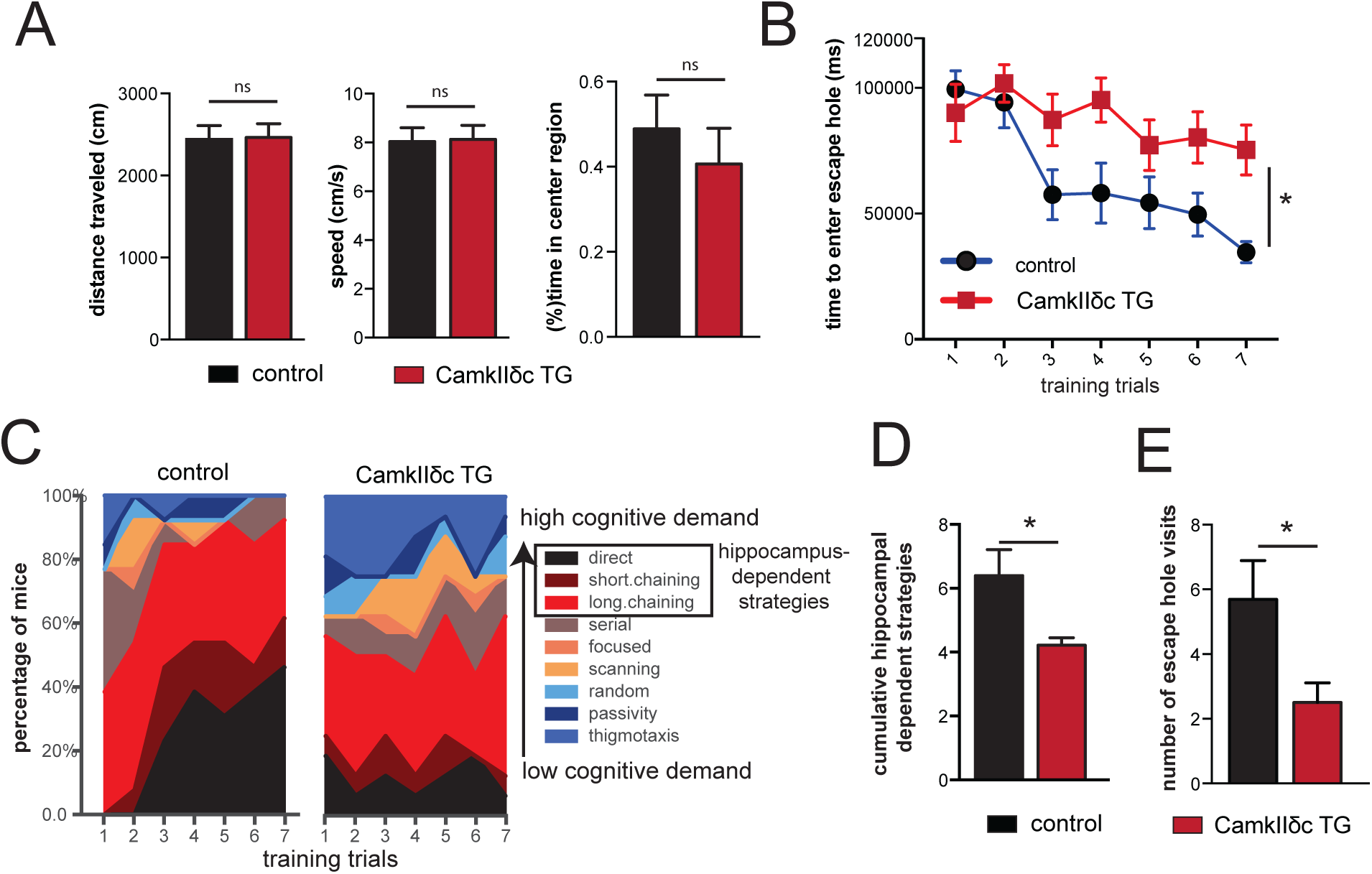
CamkIIδc TG mice display impaired hippocampus-dependent memory function. **A.** The distance traveled (left panel), the speed (middle panel) and the time spent in the center region (right panel) during a 5 min open field test was similar amongst 3 months old CamkIIδc TG (n = 16) and control mice (n = 13). **B**. The time to enter the escape hole during traning sessions of the Barnes maze test is impaired in old CamkIIδc TG (n = 16) and control mice (n = 13; two-way ANOVA, *p<0.05). **C.** Plots showing the different search strategies of CamkIIδc TG (n = 16) and control mice (n = 13) across training trials. Each strategy is labeled with a unique color. **D.** The cumulative score of hippocampus-dependent search strategies during Barnes maze training is impaired in CamkIIδc TG (n = 16) when compared to control mice (n = 13; two-tailed, unpaired t-test, *p<0.05). **E.** Number of visits to escape hole during probe test to assay memory retrieval was impaired in CamkIIδc TG (n = 16) when compared to control mice (n = 13; two-tailed, unpaired t-test, *p<0.05). Error bar indicates mean ± SEM.

### Heart failure-related down-regulation of hippocampal genes is linked to reduced neuronal H3K4-methylation

The finding that CamkIIδc mice indeed exhibit memory impairments allowed us to move on and explore our hypothesis that decreased expression of hippocampal learning and memory genes might be one of the underlying mechanisms by which heart failure leads to cognitive decline. Our gene-expression data suggest that genes down-regulated in the hippocampal CA1 region of CamkIIδc TG mice mainly reflect neuron-specific changes (See Figure 1F). In addition to pathways related to “cognition”, a major molecular process linked to these genes was protein methylation including down-regulation of H3K4-methlytransferases such as Kmt2a (see Fig 1H). Since reduced neuronal expression of Kmt2a and corresponding genome-wide reduction of H3K4me3 has been linked to memory impairment and AD (Gjoneska et al., 2015) (Kerimoglu et al., 2017), these data point to the possibility that altered H3K4-methylation may – at least in part –underlie the observed down-regulation of neuronal genes in CamkIIδc TG mice. To test this hypothesis, we retrieved and re-analyzed hippocampal RNA-seq data from mutant mice that lack the H3K4-methyltransferases Kmt2a or Kmt2b from hippocampal neurons of the adult brain and also display hippocampus-dependent memory impairment (Kerimoglu et al., 2013) (Kerimoglu et al., 2017). Our data reveal that genes decreased in CamkIIδc TG mice show a significant overlap with the genes affected in Kmt2a mutant mice (Fig 3A). In contrast, no significant overlap was seen when genes affected in CamkIIδc TG and Kmt2b mice were compared (Fig 3A). These findings are in line with the observation that Kmt2a but not Kmt2b is reduced in CamkIIδc TG mice and further supports the idea that changes in H3K4 methylation may contribute to decreased neuronal gene expression in CamkIIδc TG mice. To test this possibility directly, we decided to measure neuronal H3K4me3 in the hippocampal CA1 region of CamkIIδc TG and control mice via chromatin-immunoprecipitation followed by next-generation sequencing (ChIP-seq). Tissue of the hippocampal CA1 region was processed and subjected to FACS to isolate neuronal nuclei using an established protocol (Fig 3B)(Benito et al., 2015, Halder, Hennion et al., 2016). Afterwards H3K4me3 ChIP-seq was performed. We detected a total of 138026 H3K4me3 peaks across the entire genome. In line with previous findings from neuronal nuclei (Kerimoglu et al., 2017) and other tissues, the transcription start site (TSS) of genes was the major regulatory region where these peaks were localized (Fig 3C). When we compared H3K4me3 at the TSS of CamkIIδc TG and control mice we observed 4627 genes with decreased and 609 genes with significantly increased H3K4me3 peaks at the TSS (Fig 3D). It is important to reiterate that the ChIP-seq data stems specifically from neuronal nuclei of the hippocampal CA1 region allowing us to test the hypothesis that reduced neuronal H3K4me3 would explain the decreased expression of neuronal genes. Indeed, genes down-regulated in CamkIIδc TG (See Fig 1F, G) showed significantly reduced H3K4me3 level at their TSS (Fig 3E). In sum, these data provide strong evidence for the view that reduced neuronal H3K4me3 plays a crucial role in impaired neuronal gene expression observed in CamkIIδc TG mice and thereby contributes to heart failure induced memory loss.

**Fig. 3:**
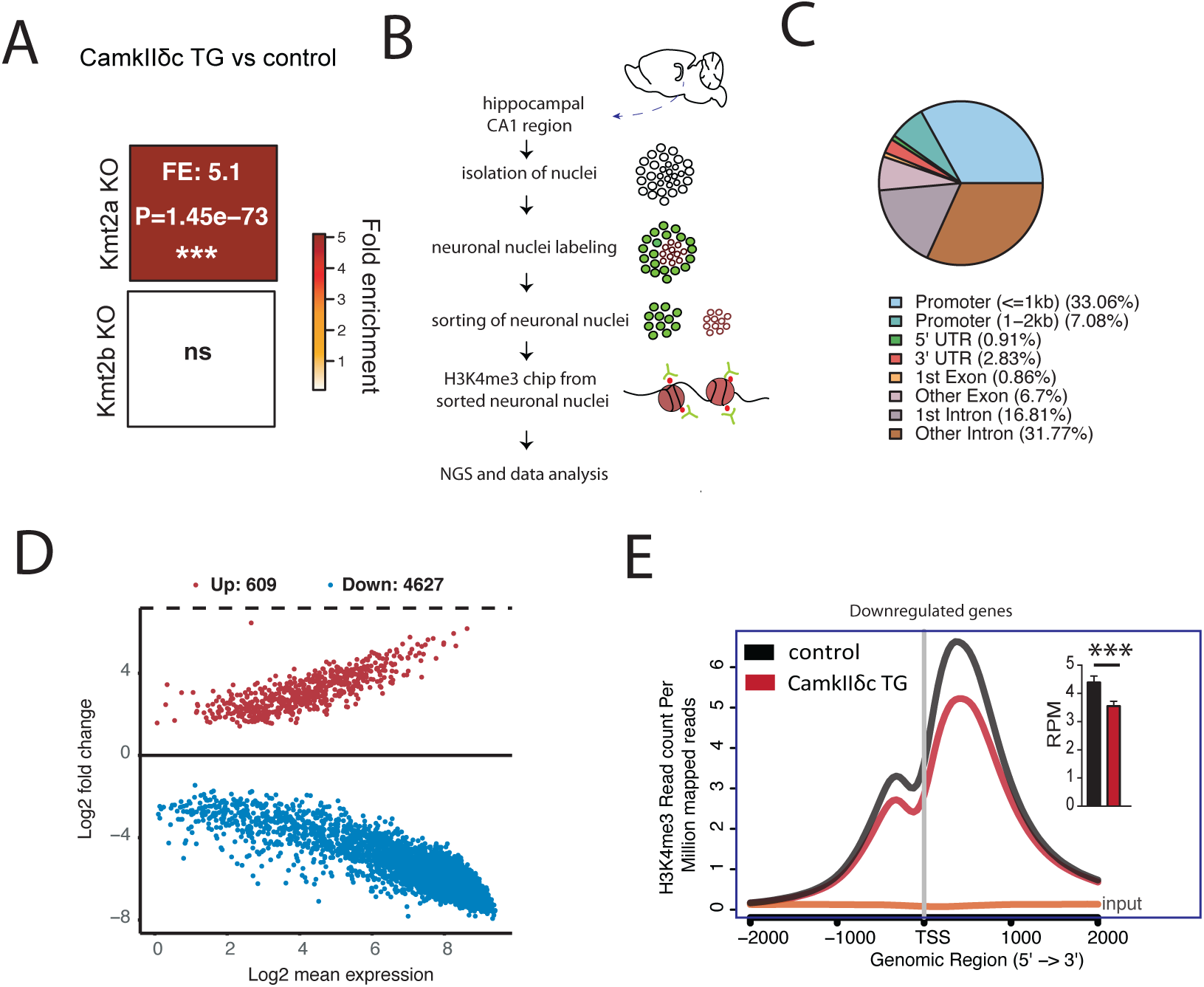
Neuronal H3K4m3 is impaired in the hippocampus of CamkIIδc TG mice. **A.** Hypergeometric overlap analysis comparing genes deregulated in CamkIIδc TG mice to genes differentially expressed in the hippocampal CA1 region of Kat2A, Kmt2A and Kmt2b knock out mice. **B.** Experimental scheme for Chip-seq analysis. **C.** Pie chart showing the distribution of H3K4me3 peaks in the neurons of the hippocampal CA1 region from CamkIIδc TG mice. **D**. MA plot showing the number of significantly altered neuronal H3K4me3 peaks when comparing CamkIIδc TG and control mice. **E.** NGS plot showing H3K4me3 peaks at the TSS of genes down-regulated in the hippocampus of CamkIIδc TG mice. Inset shows statistical analysis, (\*\*\**P* < 0.001). Error bars indicate SEM.

### Reinstating hippocampal gene-expression rescues memory impairment in CamkIIδc TG mice

Our findings point to the possibility that therapeutic strategies to increase H3K4me3 may help to ameliorate cognitive impairment in CamkIIδc TG mice and could provide a novel approach to manage cognitive impairments in heart failure patients. H3K4me3 is a chromatin mark linked to active gene-expression and euchromatin conformation. Histone-deacetylase (HDAC) inhibitors increase histone-acetylation and thereby favor euchromatin formation. Moreover, administration of HDAC inhibitors could reinstate memory function in various mouse models of neurodegenerative diseases (Fischer, 2014b) and the HDAC inhibitor Vorinostat is currently tested as therapeutic intervention in AD patients (https://clinicaltrials.gov/ct2/show/NCT03056495). Notably, HDAC inhibitors were also found to reinstate hippocampal H3K4me3 and improve spatial reference learning in mice that lack the histone-metyhltransferase Kmt2d (Bjornsson, Benjamin et al., 2014). On this basis we hypothesized that administration of Vorinostat might help to reinstate memory function in CamkIIδc TG mice, which would also provide further causal evidence for the role of altered neuronal gene-expression in HF-induced memory impairment. In a pilot experiment we found that Vorinostat was able to significantly enhance H3K9 acetylation and H3K4me3 - two euchromatin marks that are functionally related (Kerimoglu et al., 2013) (Kerimoglu et al., 2017, Stilling, Rönicke et al., 2014) - when administered to primary hippocampal neurons **(Expanded View 1)**. Thus, 2-month-old CamkIIδc TG mice were treated with Vorinostat for 1 month before behavioral testing. Another group of CamkIIδc TG mice received corresponding vehicle solution. Vehicle-treated wild type littermates served as additional control group (Fig 4A). All groups performed similarly in the open field test confirming our previous observation that CamkIIδc TG mice exhibit normal basal anxiety levels and motor function (Fig 4B). Moreover, Vorinostat had no effect on these parameters. Next, mice were subjected to the Barnes Maze paradigm to evaluate spatial reference memory. Consistent with our previous observation, vehicle treated mice displayed impaired learning behavior when compared to the corresponding wild-type group (Fig 4C). In contrast, CamkIIδc TG mice treated with Vorinostat were able to master the Barnes Maze task similar to the wild type control group (Fig 4C). Essentially, the escape latency in Vorinostat-treated CamkIIδc TG and wild-type control groups was not significantly different, suggesting that Vorinostat administration reinstates hippocampus-dependent memory function in CamkIIδc TG mice (Fig 4C). A more detailed analysis of the training procedure revealed that similar to wild type mice, Vorinostat-treated CamkIIδc TG mice eventually adopt cognitive strategies such as direct, short and long chaining strategies, while vehicle-treated CamkIIδc TG failed to do so (Fig 4D). Consistently, the cumulative cognitive score that was calculated based on these strategies (*see* methods for details) revealed a significant impairment of vehicle-treated CamkIIδc TG mice, when compared to the wild-type control group, while no such difference was observed for Vorinostat-treated CamkIIδc TG (Fig 4E). A similar observation was made when mice were subjected to the memory test after 7 training trials (Fig 4F). These data show that oral administration of Vorinostat ameliorates memory impairment in CamkIIδc TG mice.

**Fig. 4:**
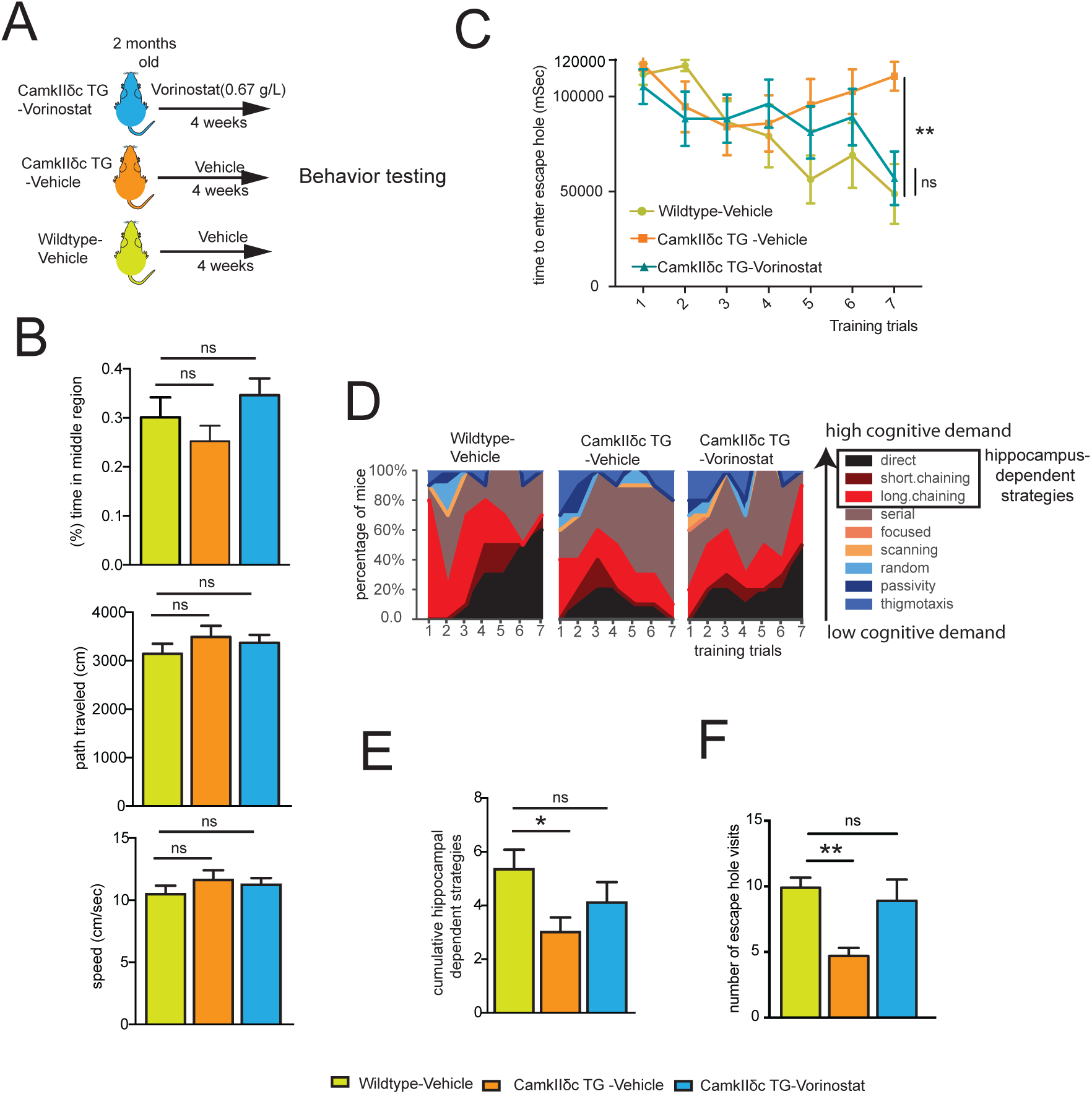
Vorinostat reinstates memory function in CamkIIδc TG mice. **A** Schematic outline of the experimental design. **B.** The distance traveled (upper panel), the speed (middle panel) and the time spent in the center region (lower panel) during a 5 min open field test were similar amongst groups (n=10/group). **C.** Latency to enter the escape hole during Barnes maze training (Two-way ANOVA, **p<0.01). **D.** Plots showing the different search strategies across training trials. Each strategy is labeled with a unique color. **E.** Cumulative hippocampus-dependent strategy scores during the Barnes maze training (One-way ANOVA, *p<0.05). F. Number of visits to the escape hole during probe test (**p<0.01). Error bars indicate mean ± SEM.

### Vorinostat ameliorates gene-expression changes in CamkIIδc TG mice

Vorinostat treatment of CamkIIδc TG mice had no significant effect on cardiac pathology (Expanded View Fig 2) suggesting that reinstatement of memory function in our experimental system is most likely linked to brain-specific processes. Thus, we analyzed gene-expression in the hippocampal CA1 region of vehicle-treated wild type mice as well as in vehicle and Vorinostat-treated CamkIIδc TG mice via RNA-seq (Fig 5A). In line with our previous observation (See Fig 1D-G), RNA-seq data analysis revealed a major deregulation of gene-expression in vehicle-treated CamkIIδc TG mice compared to the vehicle-treated wild-type control group (Expanded View Fig 3A, B). Our further analysis shows that Vorinostat could partially restore physiological gene-expression in CamkIIδc TG mice (Expanded View Fig 3B, C). The finding that Vorinostat-treatment increases the expression of genes that were down-regulated in CamkIIδc TG mice can easily be explained by the effect of Vorinostat on euchromatin formation. However, the observation that Vorinostat also decreases the expression of genes that were elevated in CamkIIδc TG mice is most likely due to additional mechanisms.

**Fig. 5:**
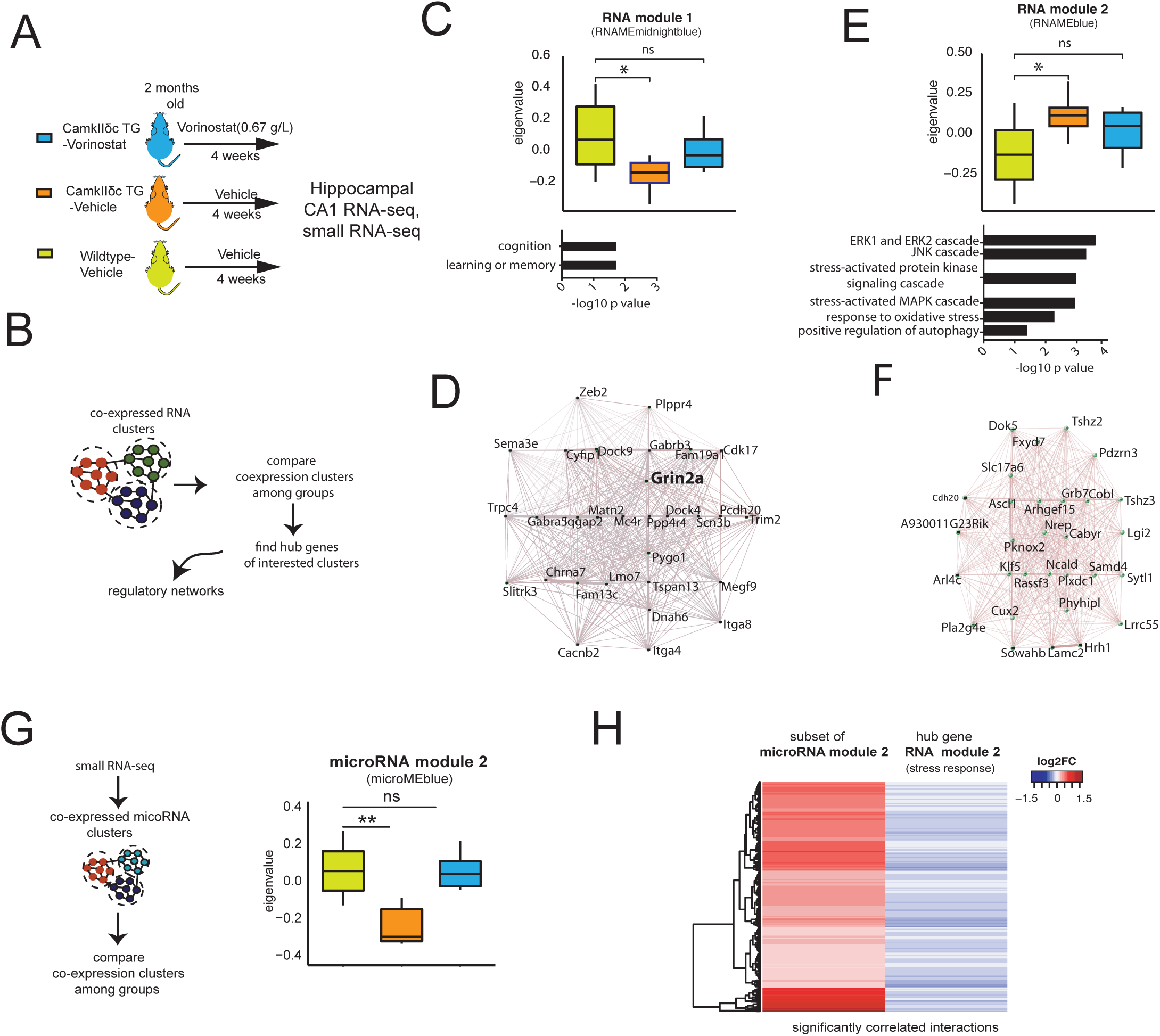
Vorinostat ameliorates pathological hippocampal gene-expression in CamkIIδc TG mice. **A.** Schematic outline of the experimental design. **B.** Scheme for WGCNA analysis. **C.** Upper panel: Expression of RNA module 1 among the three experimental groups. *p<0.05, Kruskal-Wallis test Lower panel: Gene ontology analysis of genes that are part of RNA module 1. **D.** Network representing top 30 hub genes of the gene network based on RNA module 1. **E.** Upper panel: RNA module 2 and its expression among the three experimental groups *p<0.05, Kruskal-wallis test. Lower panel: Functional annotations of the genes that are part of RNA module 2. **F.** Gene correlation network of the top hub genes (n =30) of RNA module 2. **G.** Left panel: Schematic outline of the analysis of microRNA-sequencing data. Right panel: Expression of microRNA module 2 amongst experimental groups. Kruskal-Wallis test. **p < 0.01. **H.** Heatmap showing significant negative correlation (FDR < 0.05) between microRNA members of microRNA module 2 and hub genes from RNA module 2 (see Fig 5E). Error bars indicate SEM.

To further elucidate this, we decided to investigate the RNAseq data in greater detail. Recent studies showed that the detection of regulatory co-expression modules is a suitable approach to further understand transcriptional plasticity in health and disease (Gandal, Zhang et al., 2018). To this end we performed Weighted Gene Co-expression Analysis (Langfelder & Horvath, 2008) (Fig 5B) and identified 14 different modules in the entire RNA-seq dataset (*see* methods for details). Two of these modules – namely RNA module 1 and 2 - exhibited significantly different expression amongst vehicle-treated CamkIIδc TG and wild-type control mice. RNA module 1 was decreased in vehicle-treated CamkIIδc TG mice, while its expression was partially rescued upon Vorinostat treatment (Fig 5C). Gene ontology analysis suggested that the genes of RNA module 1 are linked to cognition, learning and memory (Fig 5C). Further analysis identified a cluster of 30 hub genes within module 1. Notably, 26 of genes were shown to cause to memory impairment when their expression was manipultated (Fig 5D; supplemental table 4). In contrast, RNA module 2 was significantly increased in vehicle-treated CamkIIδc TG mice when compared to the vehicle-treated wild-type control group (Fig 5E). Expression of this cluster was partially deceased to control levels in Vorinostat-treated CamkIIδc TG mice (Fig 5E). In line with our previous analysis of up-regulated genes, the genes of RNA module 2 were mainly linked to cellular stress-related pathways (Fig 5E). In line with our previous findings the genes of RNA module 2 showed a significant overlap to genes increased in response to hypoxia in neuronal cultures (this study), human brain organoids exposed to hypoxia (Pașca, Park et al.) or ER-stress while the genes of the cognition-related RNA module 2 were decreased under the same conditions (Expanded View Fig 4).

The question remained how Vorinostat, an epigenetic drug that is linked to euchromatin formation and the activation of gene expression, would decrease the observed pathological gene-expression response linked to hypoxia and cellular stress pathways. One possible explanation is that Vorinostat induces molecular processes that antagonize this type of pathological gene expression. MicroRNAs are small non-coding RNAs that regulate cellular homeostasis via binding to a target mRNA thereby causing its degradation or inhibition of translation (Gurtan & Sharp, 2013). Compensatory microRNA responses have been described in response to various cellular stress conditions (Kagias, Nehammer et al., 2012) and we hypothesized that Vorinostat-induced microRNA expression might contribute to the therapeutic effect in CamkIIδc TG mice. To this end, we performed small RNA sequencing of the hippocampal CA1 region obtained from control and vehicle-treated CamkIIδc TG mice as well as from Vorinostat-treated CamkIIδc TG mice. Differential expression analysis revealed a number of regulated microRNAs when comparing the various conditions (supplemental table 4). To specifically identify microRNA networks that could explain the decreased expression of cellular stress-response genes upon Vorinostat treatment, we performed a weighted co-expression analysis (Langfelder & Horvath, 2008) (Fig 5G) and identified 5 microRNA modules (Expanded View Fig 5 AB). One module – namely microRNA module 2 - was significantly decreased in vehicle-treated CamkIIδc TG mice when compared to the vehicle-control group (Fig 5G), while its expression was increased to physiological levels upon Vorinostat-treatment (Fig 5G). Next, we asked whether the increased expression of microRNA module 2 would be correlated to the corresponding expression of stress-response genes increased in the hippocampal CA1 region of CamkIIδc TG mice. To this end we first performed a pairwise correlation analysis between genes and microRNAs that were differentially expressed in Vorinostat-treated vs. vehicle-treated CamkIIδc TG mice. We observed that microRNAs within microRNA module 2 showed a significant negative correlation to the hub genes of RNA module 2, representing the module linked to cellular stress responses and autophagy (Fig 5H). These data suggest that Vorinostat-treatment in CamkIIδc TG mice increases the expression of microRNAs that antagonize the expression of genes linked to pathological cellular stress. Further evidence for this view stems from the finding that these microRNAs are mainly encoded within genes that exhibit reduced hippocampal H3K4me3 in CamkIIδc TG mice (Expanded View Fig 5, C).

In sum these data suggest that aberrant neuronal gene expression plays a central role in heart failure associated cognitive decline. In turn, approaches that target these gene-expression changes could provide a novel therapeutic avenue to manage cognitive dysfunction in heart failure patients.

## Discussion

Employing a genetic mouse model for HF, we show for the first time that HF leads to substantial changes in hippocampal gene expression. The genes that were up-regulated significantly overlap with genes deregulated in neurons exposed to cellular stress such as oxidative and ER stress. These data suggest that cardiac dysfunction, which has been linked to reduced blood flow to the brain (Bikkina et al., 1994, Verdecchia et al., 2001), initiates a cellular stress response that eventually manifest at the level of neural gene-expression. Our results also reveal that these hippocampal gene-expression changes in mice suffering from HF parallel the changes observed in models for neurodegenerative diseases (Benito et al., 2015) (Gjoneska et al., 2015) (Gispert, Brehm et al., 2015) (Swarup et al., 2018). This is in line with previous reports suggesting that hypoxia-mediated oxidative and ER-stress are early and common events in neurodegenerative diseases that can trigger subsequent pathological changes associated with memory loss (Feldstein, 2012) (Xiang, Wang et al., 2017) (Butterfield & Halliwell, 2019). Indeed, further analysis of the data revealed that the hippocampal genes down-regulated in response to heart failure represent cellular processes linked to cognition and are similar to the gene-expression changes observed in models for dementia. These findings suggest that activation of cellular stress pathways might be one reason for the down-regulation of hippocampal genes essential for cognition. Support for this view stems from our observation that the sole exposure of neuronal cultures to hypoxia or ER-stress leads to the down-regulation of such neuronal gene-sets. In line with these gene-expression data we show that CamkIIδc TG mice exhibit impaired hippocampus-dependent learning and memory. Although our report of memory impairment in a heart failure mouse model is novel, these data are in agreement with various studies in humans showing that cardiac dysfunction is associated with cognitive decline and an increased dementia risk (Angermann et al., 2012) (Ampadu & Morley, 2015) (Doehner et al., 2017). Furthermore, memory impairment has been reported in animal models for acute myocardial ischemia (Evonuk, Prabhu et al., 2017) and various models for chronic cerebral hypoperfusion but the underlying molecular mechanisms remained poorly understood so far (e.g. see (Patel, Moalem et al., 2017)). How precisely activation of the various cellular stress pathways leads to the down-regulation of genes essential for cognition remains to be investigated and is likely to be multifactorial making it difficult to identify suitable targets for therapeutic intervention. From a therapeutic point of view, the fact that hippocampal genes linked to cognition are eventually decreased, might offer a more promising avenue to treat cognitive defects in HF patients, especially since these patients usually already suffer from the disease for a prolonged time period. In this context, it is important to reiterate that our data suggest that HF eventually leads to the down-regulation of gene-clusters important for cognition via processes linked to reduced histone-methylation, especially deceased levels of the euchromatin mark H3K4me3. These findings are in line with current literature showing that proper neuronal H3K4me3 is essential for memory consolidation (Gupta et al., 2010, Kerimoglu et al., 2013) (Jakovcevski et al., 2015) (Kerimoglu et al., 2017). Our data hint at a specific role of the H3K4 methyltransferase Kmt2a, which is down-regulated in the hippocampus of CamkIIδc TG mice. These findings are in line with recent reports showing that mice lacking Kmt2A in excitatory neurons of the hippocampus exhibit impaired learning and memory and decreased expression of genes implicated in cognitive function (Kerimoglu et al., 2017). Indeed, our data show that genes deregulated in the hippocampi of Kmt2a knock out mice - but not of Kmt2b - significantly overlap with deregulated genes in CamkIIδc TG mice. Taken together, these data point to a scenario in which heart failure leads to hypoxia and cellular stress, eventually driving loss of neuronal euchromatin causing decreased expression of neuronal plasticity genes essential for cognition (Fig 6). Further support for this view stems from our data that administration of an epigenetic drugs that promotes euchromatin formation reinstates memory in CamkIIδc TG mice and that this effect cannot be simply explained by improved cardiac output. These findings pave the road to a novel therapeutic approach to treat HF-induced cognitive dysfunction and lower the risk for age-associated dementia in these patients. To this end, although cerebral hypoperfusion and cellular stress appear to be initial events in the development of cognitive decline in patients suffering for cardiac dysfunction, our data suggest that they eventually lead to epigenetic changes of histone methylation in neurons. Such epigenetic alterations are known to represent long-term adaptive changes that can persist even in the absence of the initial stimulus (Fischer, 2014a). Thus, targeting the epigenome has emerged as a promising therapeutic option to treat complex and multifactorial diseases including dementia, even at an advanced stage of the disease (Fischer, 2014b) (Fischer, 2014a). In fact, previous studies showed that other risk factors for dementia such as aging (Peleg, Sananbenesi et al., 2010) (Benito et al., 2015), protein aggregation, (Kilgore, Miller et al., 2010) (Govindarajan, Agis-Balboa et al., 2011) (Benito et al., 2015) (Gjoneska et al., 2015), neuropsychiatric diseases (Nestler, Peña et al., 2015) or peripheral inflammation(Wendeln AC, Häsler LM et al., 2018) lead to similar changes representing a loss of neuronal euchromatin and reduced expression of genes linked to cognition. Of note, therapeutic strategies to reinstate euchromatin related gene-expression were able to improve memory function in such models (Benito et al., 2015) (Bahari-Javan, Varbanov et al., 2017). For example, inhibitors of histone deacetylases (HDAC) have emerged as promising candidates to treat cognitive decline, and the FDA approved HDAC inhibitor Vorinostat is currently undergoing trials in Alzheimer’s disease patients (ClinicalTrials.gov Identifier: NCT03056495). As mentioned above, oral administration of Vorinostat to CamkIIδc TG mice improved their learning and memory abilities. These findings cannot be explained by improved cardiac function, since a one-month treatment of CamkIIδc TG mice with Vorinostat had no significant effect on heart failure. However, Vorinostat treatment increased the expression of formerly downregulated hippocampal genes linked to cognition. In fact, our detailed analyses revealed that Vorinostat reinstated the expression of a specific gene cluster in which nearly every hub-gene was shown to be essential for memory function. Hence, reducing either of these genes alone was found to cause memory impairment (see supplemental table 3). While these findings are in line with the know role of Vorinostat to induced euchormatin and gene-expression, it was surprising to see that Vorinostat treated CamkIIδc TG also exhibited reduced expression of genes linked to cellular stress responses. Our data suggest that this effect is mediated via the induction of a compensatory microRNA network that downregulated cellular stress response hub genes (Fig 6). These findings are in line with the reported role of the microRNAome as one key molecular process to maintain cellular homeostasis and reports that link microRNA expression to compensatory mechanisms in various diseases (Gebert & MacRae, 2019). We cannot exclude the possibility that the improved memory function in response to Vorinostat-treatment is mediated by other mechanisms. For example, Vorinsotat also acts on non-histone proteins and has been found to suppress hypoxia signaling in cancer models (Zhang, Yang et al., 2017). Moreover, although Vorinostat increases H3K4me3, this indirect effect is most likely mediated by increased histone acetylation that generally promotes euchromatin formation. In line with this previous findings show that Kmts act in concert with histone-acetyltransferases (Kerimoglu et al., 2013) (Kerimoglu et al., 2017) (Husmann & Gozani, 2019). Nevertheless, in the future, it will be important to investigate whether therapeutic approaches that target more directly H3K4me3 are even more efficient to reinstate memory function in response to HF.

**Fig. 6.**
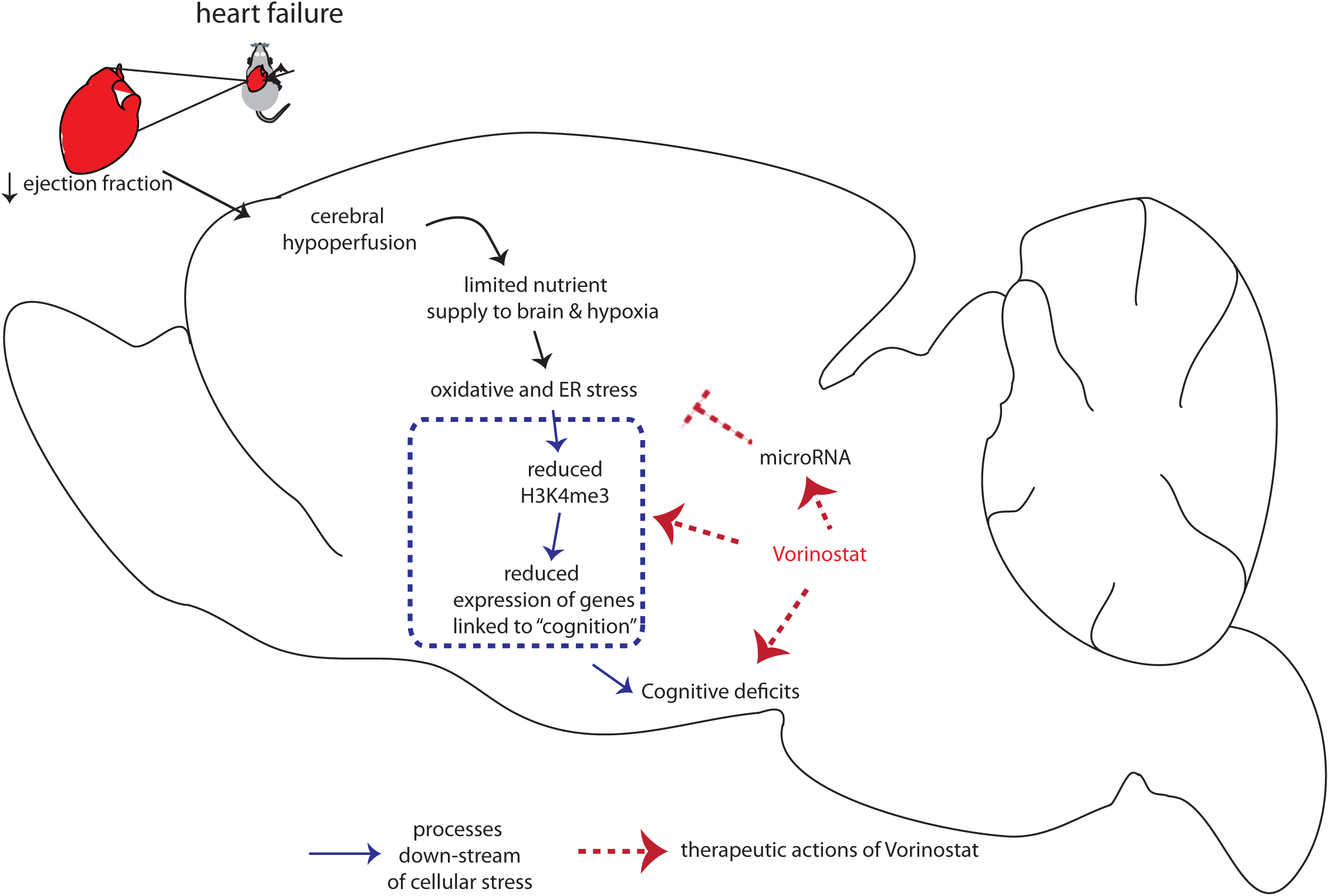
Model summarizes how heart failure contributes to memory impairment and corresponding option for therapeutic intervention. Cardiac insufficiency leads to cerebral hypoperfusion, which is in line with a hippocampal gene-expression response linked to oxidative, and ER-stress. Our data suggest that oxidative and ER-stress drive reduced expression of genes important for memory function, which involves reduced neuronal H3K4me3 representation loss of euchromatin. Administration of the HDAC inhibitor Vorinostat partially increases the expression of memory-related genes but also decreases the expression of genes linked to oxidative and ER-stress via the induction of a microRNA cluster.

In conclusion, our data elucidate the molecular mechanisms by which cardiac dysfunction contributes to cognitive impairment and suggest a key role for epigenetic neuronal gene expression. Targeting gene expression changes in the brain, through drugs such as HDAC inhibitor Vorinostat, ameliorate memory impairment and partially reinstate physiological gene expression. Thus, therapeutic strategies that target epigenetic gene expression may be a suitable approach to treat cognitive dysfunction even in chronic heart failure patients and lower their risk of developing age-associated cognitive diseases such as AD.

## Material and Methods

More detailed information is available as a supplementary material & methods file.

### Animals and tissue preparation

CamkIIδc transgenic and wild type littermates were housed in standard cages on 12h/12h light/dark cycle with food and water ad libitum. All experimental protocols were approved by a local animal care protocol. Unless otherwise stated, 3-month old mice were used for the experiments. For tissue preparation animals were sacrificed by cervical dislocation. Hippocampal sub-region CA1 was isolated, snap frozen in liquid nitrogen and stored at −80 °C. Hearts were dissected by a cut above the base of the aorta and perfused with 0.9% sodium chloride solution until blood free, snap frozen in liquid nitrogen and stored at −80°C. In addition, lung and tibia were extracted and their respective weight or length was determined.

### Echocardiography

The heart function and dimensions were examined by echocardiography using a Vevo 2100 imaging platform (Visualsonics) with 30MHz transducer (MS-400). The animals were anesthetized with isoflurane (1-2%) and M-mode sequences of the beating heart recorded in the short-axis and the long axis, respectively. The images were used to determine the left ventricular end-diastolic and end-systolic Volumes (area*length*5/6). These parameters were used to calculate the ejection fraction as indicator of left ventricular heart function. The investigator was blinded to genotype and age.

### Behaviourial tests and data analysis

#### Open Field & Barnes Maze

Open field test was performed according to a previous study (Bahari-Javan, Maddalena et al., 2012). Briefly, mice were placed gently in the middle quadrant of an open field and allowed to explore the arena for 5 minutes. The travel trajectories were recorded using VideoMot (TSE-Systems). Barnes Maze experiment was performed according to Sunyer et al (Sunyer, Patil et al., 2007).

### RNA isolation and sequencing

RNA isolation was performed using RNA Clean and Concentrator kit according to manufacturer protocol without modifications. Concentration was measured on nanodrop and quality of RNA was evaluated. For mRNA sequencing, 500 ng total RNA was used as input to prepare cDNA libraries according to Illumina Truseq and 50 bp sequencing reads were run in HiSeq 2000. For small RNA sequencing, 100 ng total RNA was used as initial input. Small RNA was enriched using size selection from based on gel. cDNA library and sequencing have been performed according to manufactureŕs protocol (NEBNext Small RNA library prep set for Illumina). Next generation sequencing was performed on HiSeq 2000 platform.

### Chromatin immunoprecipitation for H3K4me3

Chromatin immunoprecipitation was performed according to (Halder et al., 2016) with 0.2 µg chromatin and 1µg H3K4me3 (ab8580) antibody. ChIPseq library preparation was performed using NEBNext Ultra II DNA library preparation according to manufacturer’s protocol. 2nM libraries were pooled and sequenced in Illumina Hiseq 2000 with 50-bp single end reads. Details are given in Supplementary File.

### Modeling hypoxia and endoplasmic reticulum stress in primary neurons

Primary hippocampal neuronal culture was prepared as described previous (Benito et al., 2015) Experiments were performed at DIV 10. Endoplasmic reticulum stress was induced in primary hippocampal neurons using Tunicamycin (Sigma Aldrich). 2ug/mL of Tunicamycin was added to primary neuronal culture and incubated for 6 hours and compared to those treated with DMSO for same time. To model hypoxia, primary hippocampal neuronal cultures were incubated in normoxia (20% O_2_) in a standard cell culture incubator for 10 days before they were used in an experiment. For hypoxic conditions cells were incubated at 1% O_2_ for 4 hours using the *in invivo_2_* 400 hypoxia workstation (Baker Ruskin). Cells from the same isolation were kept in normoxia at 20% O_2_ as a control.

### Quantitative RT-PCR

qPCR (q-PCR) primers were designed using Universal probe library Assay Design Center and were purchased from Sigma. Transcriptor High Fidelity cDNA Synthesis Kit (Roche) was used to prepare cDNA. UPL probes were used for quantification and data was normalized to HPRT1 expression as internal control. Relative gene expression was analyzed by 2−ddCt method. Primer sequences are summarized in Supplementary File.

### Western blot

Western blot was performed according to previous study (Bahari-Javan et al., 2012). To quantify H3K4me3 and H3K9ac levels H3K4me3 (abcam, ab8580) antibody, H3K9ac (abcam, ab4441) antibody were used respectively. Unmodified H3 level measured with H3 antibody (abcam, ab1791) was used as internal control.

### Bioinformatics analysis

Bulk RNA Sequencing data analysis has been performed according to (Benito et al., 2015). Small RNA sequencing files were analyzed according to using miRDeep2. A wrapper of the steps applied during mapping and counting is available as package (https://github.com/mdrezaulislam/MicroRNA). Differentially expressed genes or microRNAs were determined using mixed linear model accounting for technical and biological covariates.

For linear mixed effects model, limma in R was implemented. Biological processes were analyzed using Gene Ontology (http://geneontology.org/). For pathway analysis KEGG (https://www.genome.jp/kegg/), Reactome (https://reactome.org/) databases were used. Exon counts were generated using DEXSeq (http://bioconductor.org/packages/DEXSeq/). Hypergeometric test analysis was performed using GeneOverlap (http://bioconductor.org/packages/GeneOverlap/). H3K4me3 peaks were called using MACS2. Chip peaks were mapped at promoter (TSS ± 2kb) of genes using ngs.plot. Weighted co-expression analysis for both microRNAs and mRNAs was performed using WGCNA (Langfelder & Horvath, 2008). External gene expression datasets that have been used from other studies were downloaded from NCBI GEO (https://www.ncbi.nlm.nih.gov/geo/) and mapped in house to have consistency in results. Details of microRNA promoter and host genes and other analysis are summarized in Supplementary File.

### Statistical analysis

All the statistical analyses as mentioned in the main text are performed in Prism (version 7.0) or in R.

### Accession number for RNA-seq and ChIP-seq data

Raw data for all next-generation sequencing samples can be accesses via the following accession number via GEO database: RNA-seq (GEO*), smallRNA-seq (GEO*), ChIP-seq (GEO*) *will be made available upon acceptance of the manuscript.

## Acknowledgments

The authors thank Susanne Burkhard, Alessya Kretzschmar and Sabrina Koszewa for technical support. This work was supported by the funds from the SFB1002 (D04) of the German Research Foundation (DFG) to KT, DMK was supported by funds from the IRTG1816 and DFG Ka1269/13-1 of ther DFG. AF was supported by the DZNE, the DFG under Germany’s Excellence Strategy - EXC 2067/1 390729940 and the Hans and Ilse Breuer Foundation.

## Authors contribution

MRI coordinated the project, performed experiment, analyzed data, and wrote the manuscript. DL breed mice and performed echo, MSS performed ChIP-seq, RMH contributed to some behavioral experiments and echo analysis; TB performed tissue dissection, MSS performed qPCR analysis, JC performed immunoblot analysis, EV contributed to the analysis of Barnes Maze data, CS, DMK and AZ performed hypoxia experiments, FS, KT, AF designed the project, KT and AF wrote the manuscript and co-supervised all work related to this manuscript.

---

**Expanded view Fig 1.**
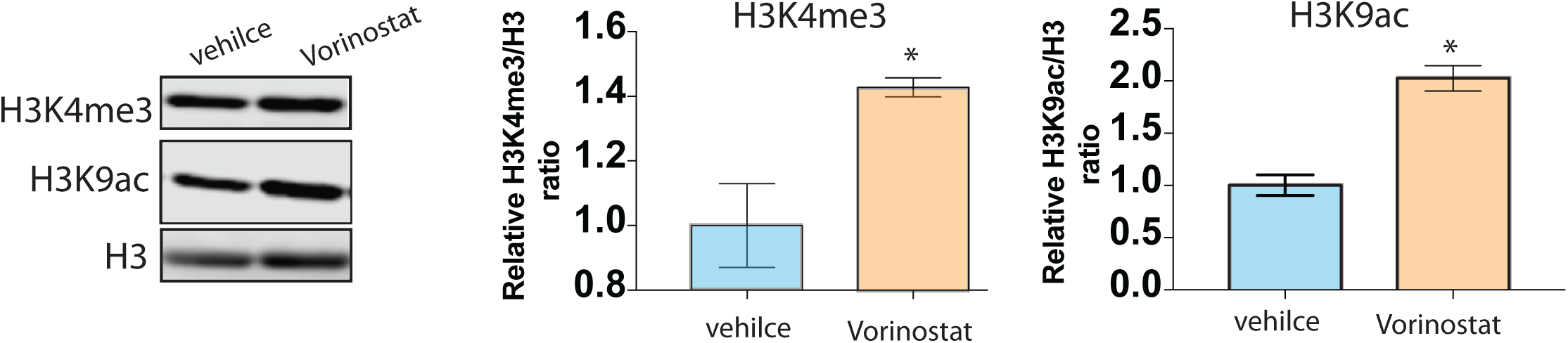
Vorinostat increases H3K4me3 level in primary neurons. Primary mouse hippocampal neuronal cultures (10 DIV; n=3/group) were treated for 1h with Vorinostat (1µm) or vehicle before proteins were isolated and subjected to immunoblot analysis. Left panel: Representative immubnoblot analysis. H3 was used as loading control. Right panels: Semi-quantitative analysis of immunblot analysis from two independent experiments (n=3/group and experiment). Relative Intensity was normalized to H3 level. The data reveal that Vorinostat treatment significantly increases bulk levels of H3K4me3 and H3K9ac. *P < 0.05, t-test; Error bars indicate SEM.

**Expanded View Fig 2:**
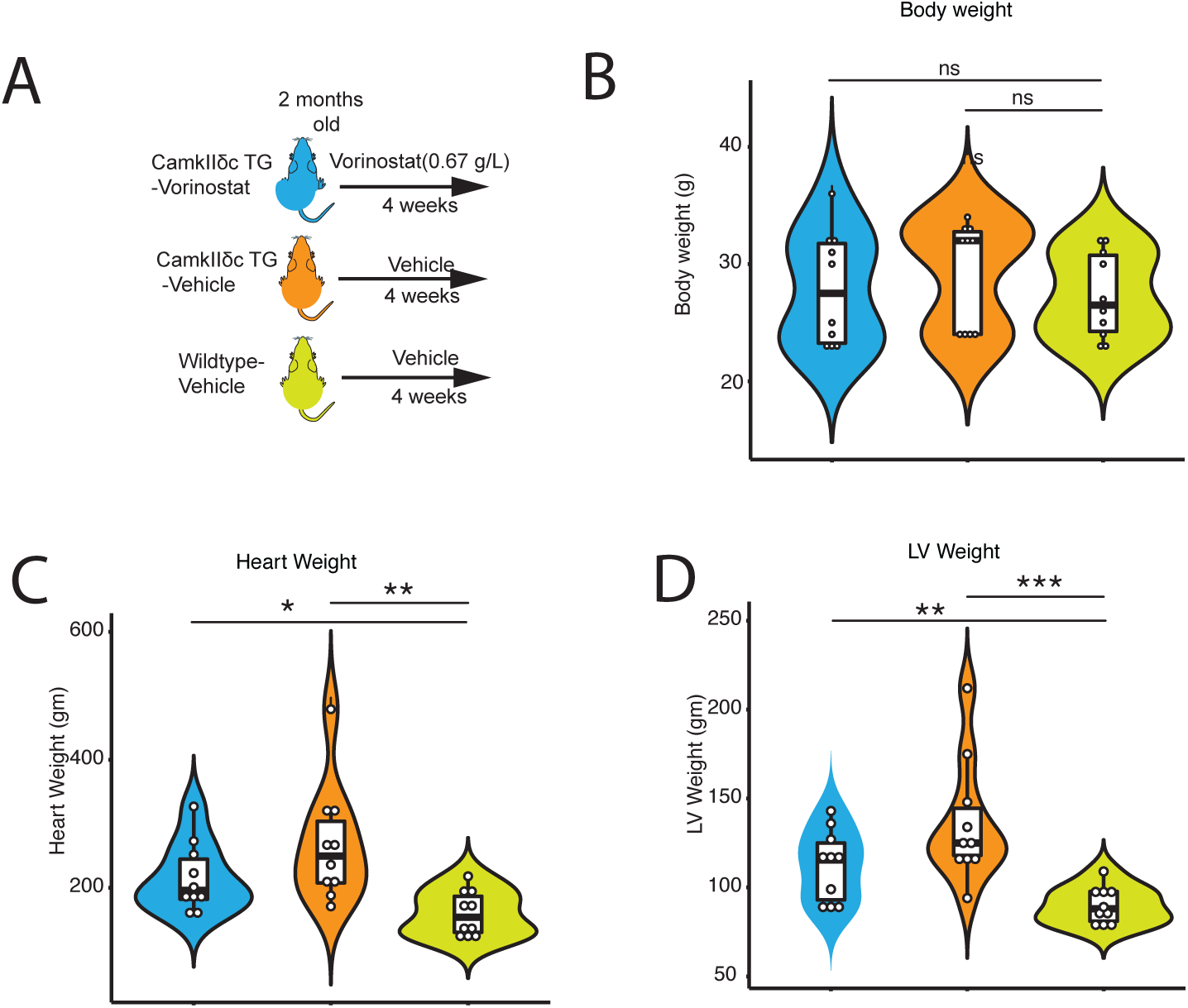
Cardiac function is not significantly affected in CamkIIδc TG mice upon Vorinostat treatment. **A.** Scheme of the experimental design. Vorinostat treat was initiated at 2 month of age and analysis was performed at 3 month of age. n= 10/group. **B.** Violin plot showing that body weight was similar amongst groups. **C.** Violin plots showing that heart weight and left ventricle (LV) weight **(D)** was increased in vehicle and Vorinostat-treated CamkIIδc TG mice when compared to the wild-type vehicle control group. ns; not significant, *P< 0.05, **P < 0.01, *** P< 0.001, Kruskal wallis t-Test.

**Expanded View Fig 3:**
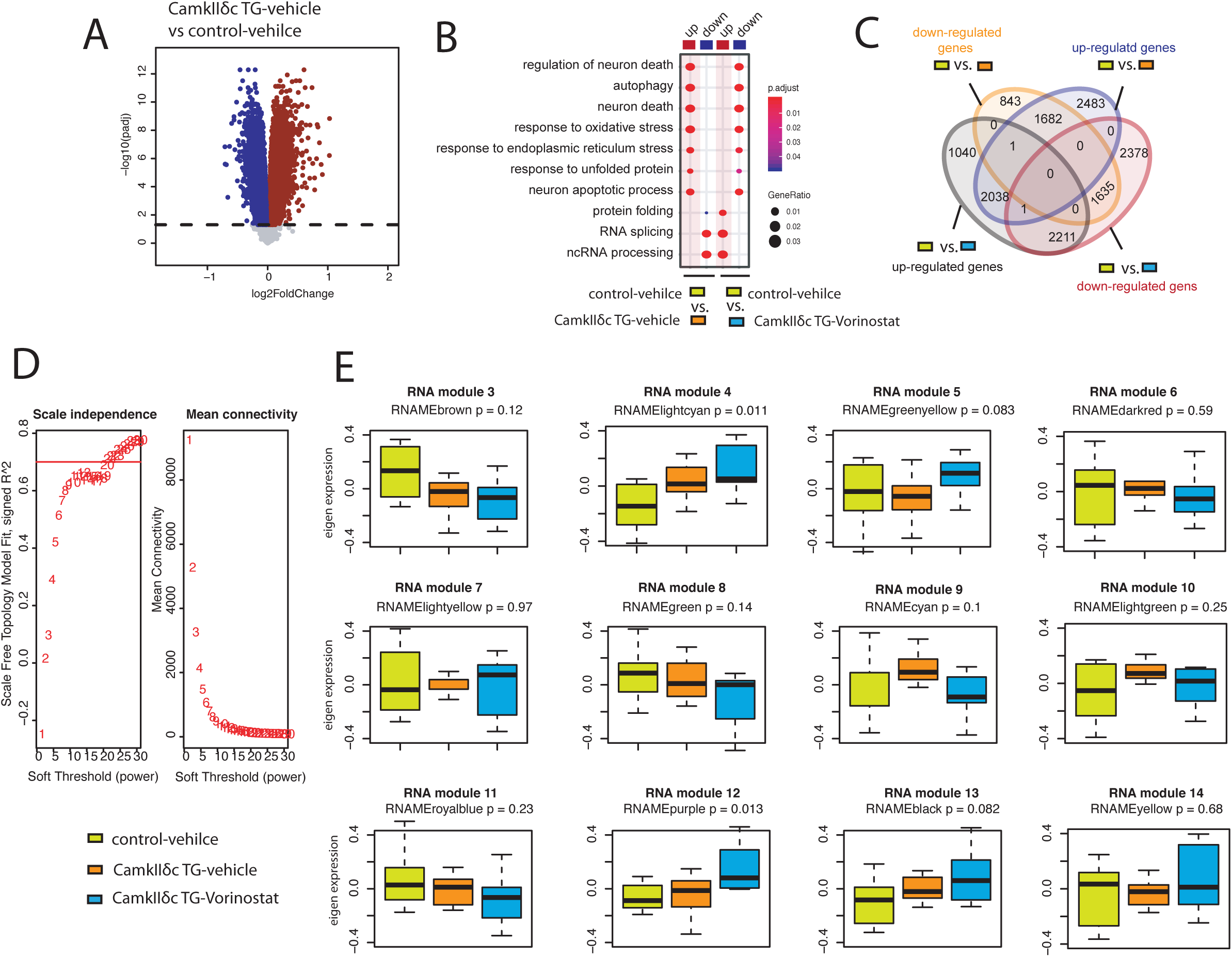
Weighted gene co-expression analysis upon Vorinostat treatment. **A.** Volcano plot showing significantly deregulated genes in the hippocampal CA1 region, when comparing vehicle-treated CamkIIδc Tg mice to vehicle-treated control mice (FDR < 0.05). Up- and down-regulated genes are represented in darkred and darkblue colors respectively. **B.** Pathways affected in the hippocampal CA1 region when comparing vehilce-treated wild type vs.CamkIIδc mice to Vorinostat-treated CamkIIδc vs vehicle-treated wild type mice. Note that Vorinostat-treatment ameliorates pathways affected in CamkIIδc TG mice for pathway increased and decreased under pathological conditions. **C.** Venn diagram showing common and uniquely deregulated genes between groups. **D.** Soft power selection based on scale independence and mean connectivity for different modules identification in WGCNA. E. Different modules representing distinct expression patterns among groups. Y axis representing eigen expression of given cluster/module.

**Expanded View Fig. 4:**
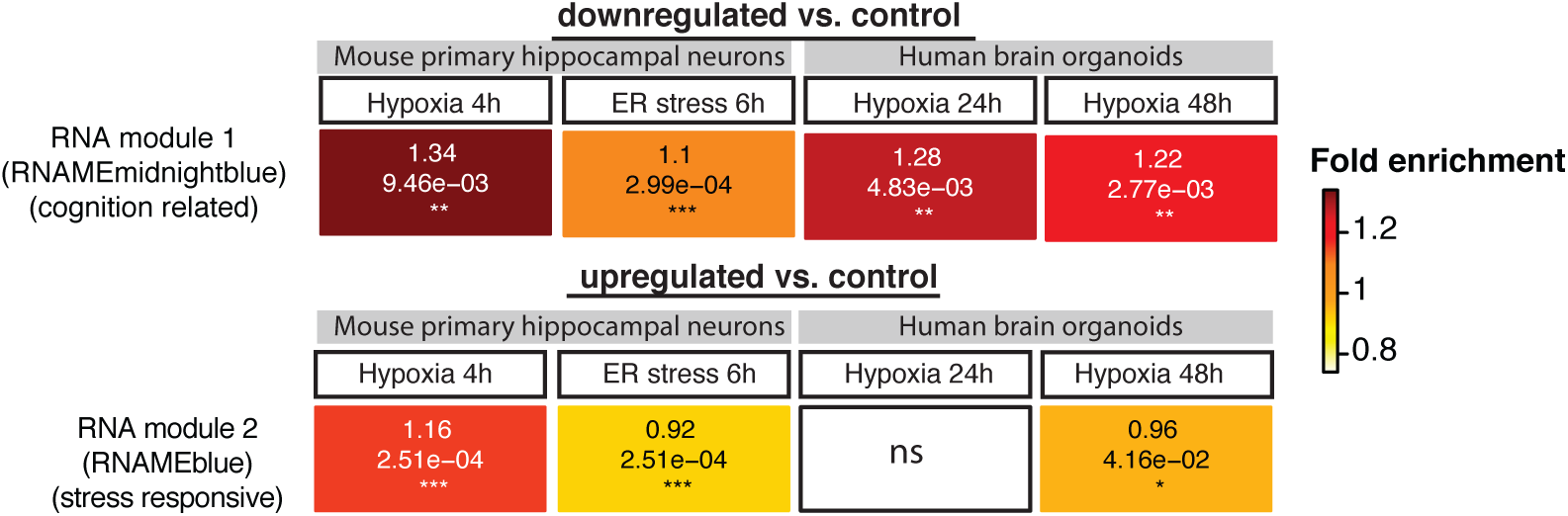
Hypogeometric overlap analysis comparing conserved the gene-expression networks RNA module 1 and 2 to hypoxic and ER stress conditions. Heatmaps summarizing results from hypergeometric tests for genes in RNA module 1 and 2 with stress conditions in different experimental settings. Hypoxia (1% O2, 4h) and endoplasmic stress (tunicamycin 2 ug/mL, 6h) was modeled in primary hippocampal neurons. Gene expression data on hypoxia from human brain organoid data is retrieved from Pasca et al, 2019. Up and down regulated genes (FDR<0.05) were determined by comparing to corresponding controls. Enrichment significance cutoff: FDR < 0.05. Color intensity represents fold enrichment.

**Expanded View Fig 5.**
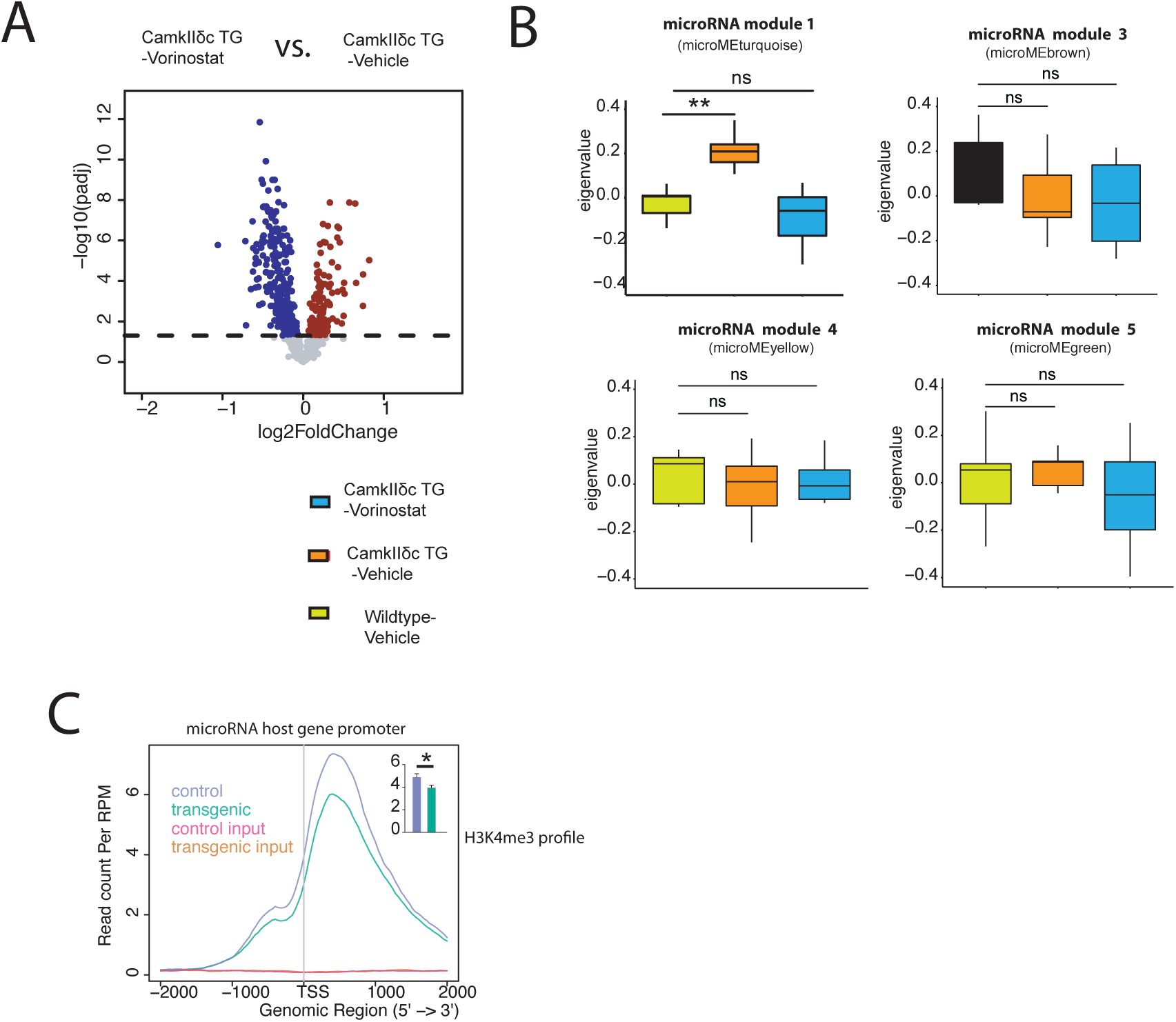
Vorinostat induced microRNA expression changes in the hippocampal CA1 region of CamkIIδc TG mice. **A**. Volcano plot showing differentially expressed microRNAs when comparing Vorinostat-treated to vehicle treated CamkIIδc TG mice. **B.** Expression of microRNA modules from WGCNA analysis in three experimental groups. wiltd type vehicle: n = 9; CamkIIδc TG-vehile n = 7; CamkIIδc TG-Vorinostat n = 10; kruskal wallis test. **C.** H3K4me3 profile at promoter of genes that harbor microRNAs. H3K4me3 level at promoter of these genes is signifcantly reduced in transgenic mice. Unpaired t-test, *p<0.05. Error bars indicate SEM.

